# The stoichiometry of minor-to-major pilins regulates the dynamic activity of the type IVa competence pilus in *Vibrio cholerae*

**DOI:** 10.64898/2026.01.17.700090

**Authors:** Nicholas D. Christman, Triana N. Dalia, Jennifer L. Chlebek, Ankur B. Dalia

## Abstract

Type IVa pili (T4aP) are bacterial surface appendages that perform various functions including twitching motility, reversible surface attachment, microcolony formation, surface sensing, and DNA uptake for natural transformation. Pivotal to each of these functions is the ability of T4aP to be dynamically extended and retracted from the cell surface. However, the factors that regulate this dynamic activity remain poorly understood in most systems. To address this question, we employ the competence T4aP from *Vibrio cholerae* as a model system. T4aP are composed of major and minor pilin subunits, named based on their relative abundance in the pilus filament. Prior work has established that minor pilins form a complex that initiates T4aP assembly. This allows for the subsequent addition of major pilins to the filament, which promotes T4aP extension. Here, we uncover that the stoichiometry of minor-to-major pilins is a crucial determinant of T4aP dynamic activity. Specifically, we show that either 1) overexpressing minor pilins or 2) underexpressing the major pilin results in a dramatic increase in the frequency of T4aP dynamics. These results indicate that the stoichiometry of minor-to-major pilins, not their absolute abundance, is one mechanism that regulates T4aP dynamic activity.

**AUTHOR SUMMARY:** Type IVa pili (T4aP) are a broadly conserved family of filamentous bacterial appendages that help bacteria colonize surfaces, move towards or away from stimuli, and gain new traits through a mechanism of horizontal gene transfer called natural transformation. T4aP are primarily composed of protein subunits called major and minor pilins, named based on their relative abundance in the pilus filament. Bacteria can dynamically extend and retract pilus filaments from their surface through polymerization and depolymerization of these pilins. This dynamic activity is critical for the activities that T4aP carry out. However, the factors that regulate this dynamic activity remain incompletely understood. Here, we find that the ratio of minor-to-major pilins is one factor that regulates the frequency of dynamic activity. Minor pilins are a universally conserved feature of T4aP in diderms. So, it is tempting to speculate that the minor-to-major pilin ratio represents a broadly conserved mechanism for controlling dynamic T4aP activity in diverse bacterial species.

## INTRODUCTION

Type IV pili (T4P) are dynamic filamentous nanomachines found in diverse bacteria and archaea and are subdivided into several different classes. Bacterial T4P are members of one of four classes: T4aP (*e*.*g*., the *Vibrio cholerae* competence T4P), T4bP (*e*.*g*., the *V. cholerae* toxin co-regulated pilus), T4cP (*e*.*g*., the *Caulobacter crescentus* tad pilus), and T4dP (*e*.*g*., the *Streptococcus pneumoniae* Com pilus). This publication focuses on T4aP, which represents the most thoroughly characterized T4P system. T4aP filaments are primarily composed of a repeating protein subunit known as the major pilin [1-3]. However, T4aP filaments also include additional, low abundance pilin proteins known as minor pilins that are crucial for T4aP assembly and function.

Minor pilins play at least two roles in T4aP biology: 1) initiating pilus assembly and 2) mediating the activity of the pilus. A subset of “core” minor pilins (commonly 4 distinct pilins) form a complex that primes the T4aP machines of diderms for filament assembly, and without them, pili are rarely assembled [4-9]. Notably, the complex of minor pilins that prime T4aP structurally resemble the minor pilins that prime assembly of the type II secretion system (T2SS) and T4dP [10-15]. Because the minor pilin complex primes the T4aP machine [4, 6] for the subsequent addition of major pilin subunits at the base of the filament, the complex of core minor pilins is likely localized at the tip of extended pili [4, 6, 12, 16, 17]. Minor pilins have been shown to mediate many of the functions of T4aP and T4dP, namely reversible surface attachment, twitching motility, microcolony formation, surface sensing, and DNA-binding for natural transformation [10-12, 16-27]. These activities can be attributed to core and auxiliary minor pilins, depending on the T4aP system. In addition to the tip-associated minor pilin complex, minor pilins may also be inserted along the length of pili [7, 28].

Critical to T4aP function is the ability of individual T4aP machines to actively extend and retract a pilus, a property we define as the “frequency of dynamic activity” (*i*.*e*. how often a pilus is extended and/or retracted from the cell surface) [29]. While dynamic activity is a necessary property of T4aP, not all systems exhibit the same frequency of dynamic activity [16, 30-36]. One possibility is that the frequency of dynamic activity is optimized for the function that each T4aP carries out. For example, twitching motility may require numerous dynamic T4aP per cell to promote coordinated movement, while DNA uptake for natural transformation can feasibly occur via the action of a single dynamic T4aP machine. Despite dynamic pilus extension / retraction being crucial for T4aP function, the factors that regulate the frequency of dynamic activity remain poorly understood in most systems. Here, we use the competence T4aP from *V. cholerae*, an emerging model system for studying T4aP dynamics [16, 30, 37-43], to address this question.

## RESULTS

### Overexpression of minor pilins increases the frequency of T4aP dynamic activity

*V. cholerae* competence T4aP dynamics can be directly observed by epifluorescence timelapse microscopy using strains where the major pilin, *pilA*, contains a cysteine knock-in (*pilA*^*S67C*^) that allows for labeling pili with an Alexa Fluor 488 conjugated maleimide dye (AF488-mal) [16, 40]. In the parent strain, competence T4aP are rapidly extended and retracted; and the majority of cells exhibit ≤2 dynamic pili during a 2 min timelapse (a dynamic event is defined as any pilus that is extended, retracted, or extended and retracted during the 2 min timelapse; **Fig. 1, 2A-C**) [16, 43]. However, each cell has the capacity to assemble many more pili. This is supported, in part, by the observation that deletion of the retraction motor ATPase, *pilT*, results in hyperpiliated cells (**Fig. 1**) [16, 30, 37, 38, 42]. This suggests that the frequency of T4aP dynamic activity is constrained in the parent.

**Fig. 1.**
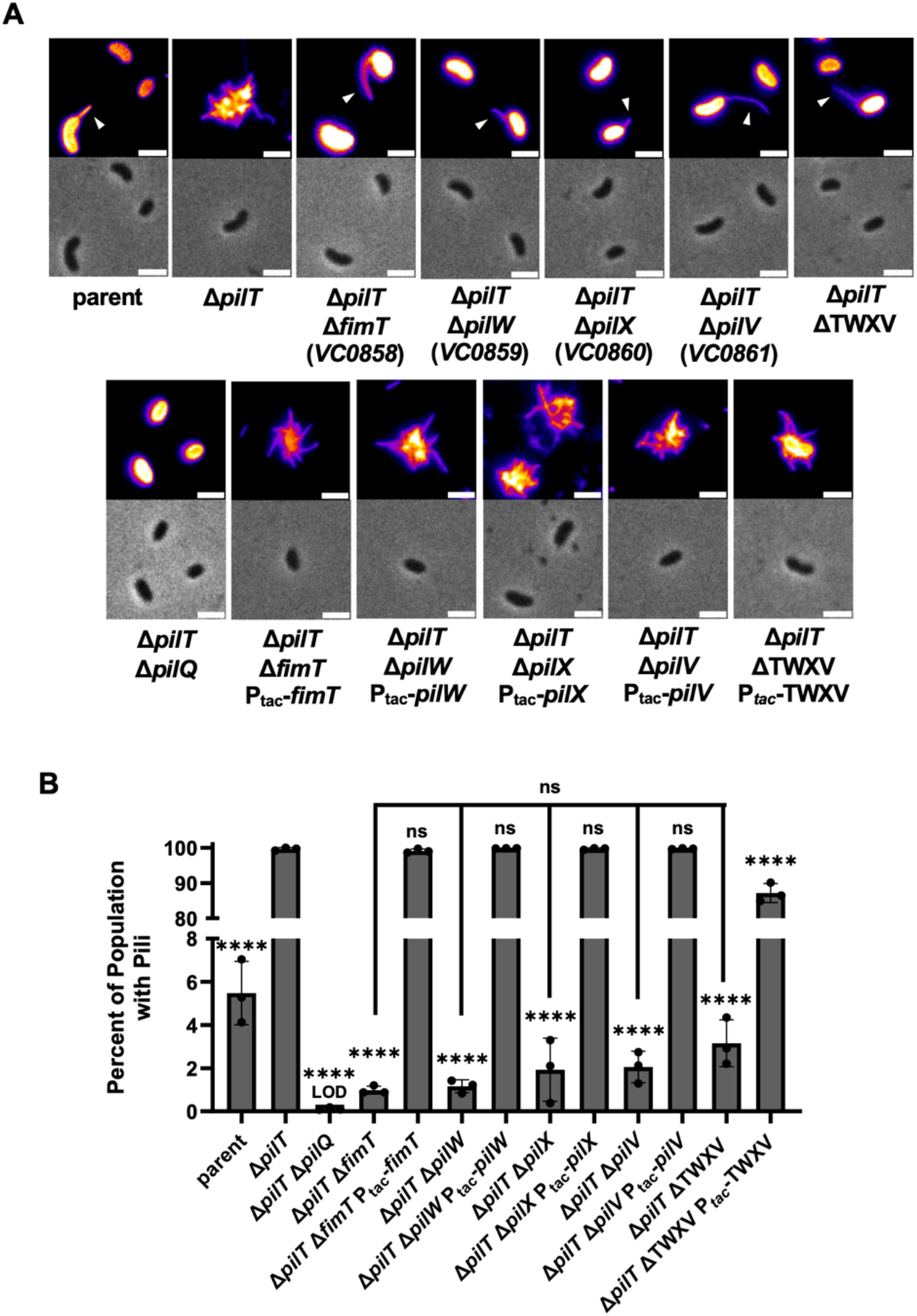
Empirical validation of the core competence T4aP minor pilins in *V. cholerae*. (**A**) Images of a representative piliated cell of the indicated strains stained with AF488-mal. All strains were induced with 100 µM IPTG. Images are false colored with the “Fire” LUT in Fiji to make rare pili easier to see (white arrows). The Δ*pilT* Δ*pilQ* strain lacks surface pili, so only unpiliated cells are shown. Scale bar, 2 µm. (**B**) Quantification of surface piliation in the indicated strains stained with AF488-mal, as depicted in **A**. Data in **B** is from 3 independent biological replicates and shown as the mean ± SD. The number of cells quantified varied for each strain and replicate. The total number of cells analyzed for each strain: parent = 1496, Δ*pilT* = 2478, Δ*pilT* Δ*pilQ* = 2917, Δ*pilT* Δ*fimT* = 4003, Δ*pilT* Δ*pilW* = 3331, Δ*pilT* Δ*pilX* = 3791, Δ*pilT* Δ*pilV* = 3233, Δ*pilT* Δ*fimT* Ptac-*fimT* = 2483, Δ*pilT* Δ*pilW* P_tac_-*pilW* = 2489, Δ*pilT* Δ*pilX* P_tac_-*pilX* = 2560, Δ*pilT* Δ*pilV* P_tac_-*pilV* = 3011, Δ*pilT* ΔTWXV = 6585, Δ*pilT* ΔTWXV P_tac_-TWXV = 4860. Statistical comparisons were made by one-way ANOVA with Tukey’s multiple comparison test of the log-transformed data. NS, no significance; **** = *p* < 0.0001. LOD, limit of detection. Statistical identifiers directly above bars represent comparisons to Δ*pilT*.

**Fig. 2.**
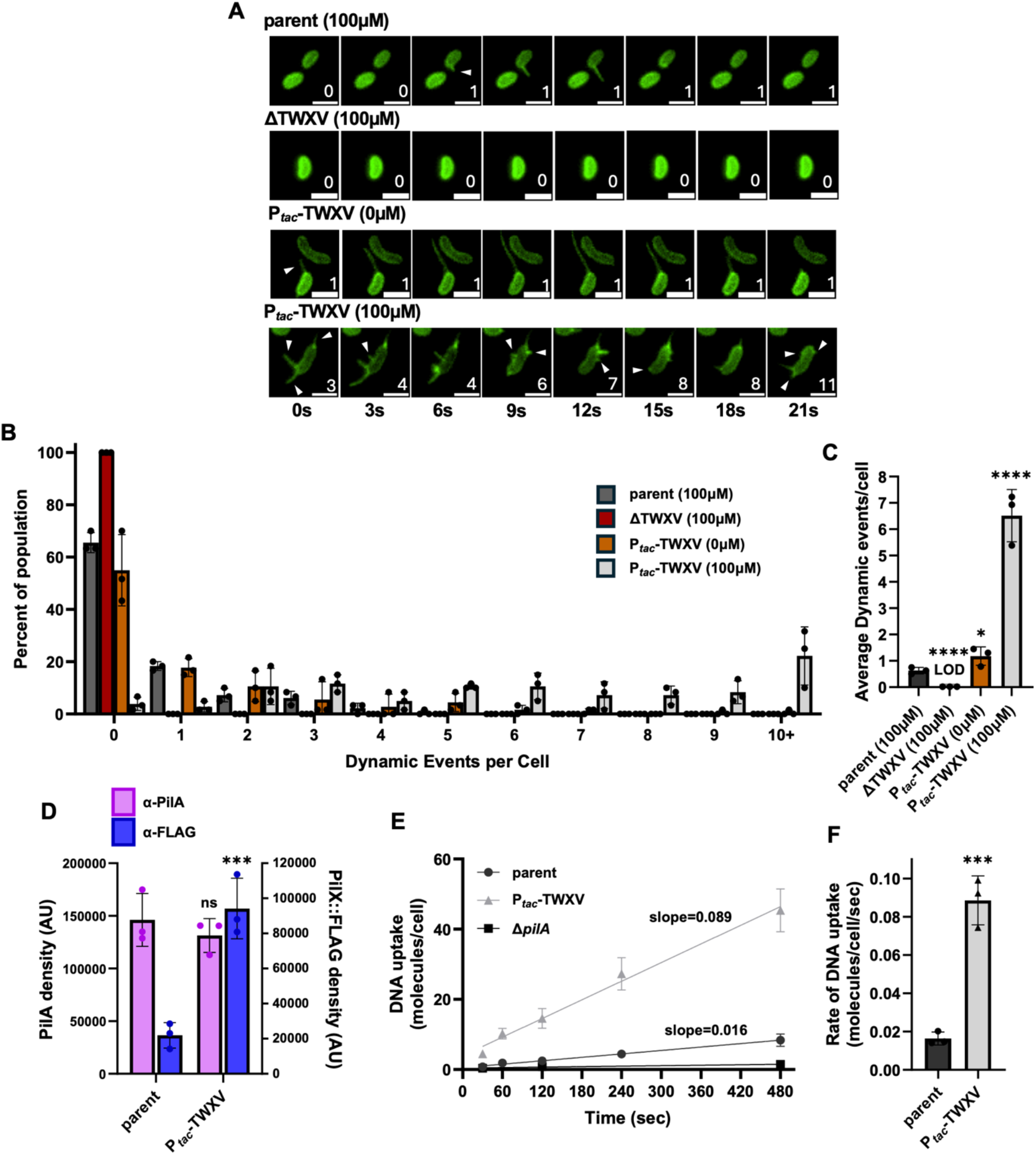
Overexpression of minor pilins increases the frequency of T4aP dynamic activity. (**A**) Representative montages of timelapse imaging of AF488-mal labeled cells from the indicated strains. Cells were induced with the indicated concentration of IPTG. Each dynamic event is demarcated by a white arrow. Additionally, a running tally of the total dynamic events in each montage is included in the lower right corner. Scale bar, 2 µm. The frequency of T4aP dynamics was quantified from timelapse imaging of AF488-mal labeled cells (as depicted in **A**) and the (**B**) distribution and (**C**) average dynamics per cell are plotted. For **D**-**F**, all strains were induced with 100 µM IPTG. (**D**) Quantification of PilA and PilX::FLAG via semi-quantitative western blots of the indicated strains (see **Fig. S4** for standard curves and details). For western blots, the parent strain contains an internally FLAG-tagged allele of *pilX* at the native locus = *pilX*::FLAG; and the P_tac_-TWXV strain had the same *pilX*::FLAG allele at both the native locus and in the ectopic construct = *pilX*::FLAG P_tac_-TWX(::FLAG)V. Blots were stained with either α-PilA (purple bars) or α-FLAG (blue bars) primary antibodies. (**E**) DNA uptake into the periplasm of the indicated strains induced with the indicated concentration of IPTG was kinetically monitored by qPCR. Lines of best fit were determined by linear regression. The slope of each line is shown, which represents the rate of DNA uptake. (**F**) The rate of DNA uptake of the indicated strains was derived from the slope of the plots depicted in **E**. Data from **B-F** are from 3 independent biological replicates and shown as the mean ± SD. Statistical comparisons in **C** were made by one-way ANOVA with Tukey’s multiple comparison test of the log-transformed data and in **D** and **F** were made by unpaired Student’s t-test of the log-transformed data. NS, no significance; * = p < 0.05, ** = *p* < 0.01, *** = *p* < 0.001, **** = *p* < 0.0001. LOD, limit of detection. Statistical identifiers directly above bars represent comparisons to the parent.

Because core minor pilins are essential for initiating pilus assembly, we hypothesized that a low abundance of core minor pilins may be one factor that constrains the frequency of dynamic activity (**Fig. 2A-C**). To test this, we first needed to establish the identity of the core pilins of the *V. cholerae* competence T4aP. It was previously hypothesized that VC0858, VC0859, VC0860, and VC0861 encode the competence T4aP minor pilins [42]. Furthermore, recent work has established that diverse members of the type IV filament family contain a structurally conserved core of 4 minor pilins that initiate pilus assembly [4, 6, 10, 12, 13]. AlphaFold modeling suggests that the minor pilins encoded by VC0858-VC0861 form this initiation complex (**Fig. S1**). To reflect this conservation and to ease comparisons between T4aP systems, we named these putative minor pilins based on the convention established in *Myxococcus xanthus* and *Pseudomonas aeruginosa*: VC0858 = *fimT*, VC0859 = *pilW*, VC0860 = *pilX*, and VC0861 = *pilV*.

Because core minor pilins prime T4aP extension, deletion of minor pilin genes results in a severe reduction in pilus assembly. Pilus assembly in minor pilin mutants, however, is not completely eliminated [4, 7, 8, 44, 45], which may be due to cross-complementation via the minor pilins of the T2SS [4, 5], which form a complex that is structurally homologous to the T4aP minor pilins (**Fig. S1**) [5, 13, 14]. Thus, we mutated each of the four putative competence T4aP minor pilin genes to test whether they exhibited the characteristic phenotype of reduced, but not eliminated, surface piliation. *V. cholerae* competence T4aP are highly dynamic structures [16, 40]. This reduces the appearance of surface pili in static images, which limits the dynamic range for assessing T4aP assembly. Specifically, in the parent, only ∼6% of cells have an extended pilus in a static image (**Fig. 1B**) despite >40% of cells assembling a pilus within a 2 min timelapse (**Fig. 2A-C**). Thus, to sensitize these experiments to more rigorously assess pilus assembly, we utilized a Δ*pilT* background (*i*.*e*., assembled pili will remain static on the cell surface in the Δ*pilT* background) where >99% of cells have surface pili in static images (**Fig. 1**). Mutating *fimT, pilW, pilX*, and *pilV* individually or altogether (ΔTWXV), severely reduced, but did not eliminate, surface piliation (**Fig. 1**), which is consistent with them encoding the core minor pilins. By contrast, mutating the essential competence T4aP machine component *pilQ* [42, 46] results in a complete loss in surface piliation (**Fig. 1**). Importantly, the surface piliation of each minor pilin mutant could be complemented (**Fig. 1**). Also, deletion of all 4 minor pilins (ΔTWXV) ablates observable T4aP dynamic activity (**Fig. 2A-C**), which is consistent with their critical role in promoting efficient competence T4aP assembly.

As mentioned above, the rare piliation observed in Δ*pilT* ΔTWXV mutants may be due to T2SS minor pilins, which have been shown to cross-complement and promote T4aP assembly in *P. aeruginosa* and *E. coli* [4, 5]. Consistent with this, we found that deletion of both the competence T4aP minor pilins and T2SS minor pilins ablates surface piliation (ΔHIJK ΔTWXV Δ*pilT*; **Fig. S2**). Furthermore, we found that ectopic expression of T2SS minor pilins in Δ*pilT* ΔTWXV increases surface piliation (P_*tac*_-HIJK ΔTWXV Δ*pilT*; **Fig. S2**). Together, these results strongly suggest that the T2SS minor pilins can cross-complement and promote assembly of the *V. cholerae* competence T4aP.

Next, we sought to test whether the abundance of the competence T4aP minor pilins regulates dynamic activity. We hypothesized that if low expression of minor pilins constrains dynamic activity, then overexpression of these minor pilins should markedly increase the frequency of T4aP dynamics. The native *fimT, pilW, pilX*, and *pilV* genes are all expressed in an operon; and these genes are partially overlapping, which indicates that they are likely translationally coupled [47, 48] to reinforce expression at a fixed stoichiometry. To overexpress these core minor pilins while preserving their translational coupling, we cloned the entire *fim<u>T</u>*-*pil<u>W</u>*-*pil<u>X</u>*-*pil<u>V</u>* operon (TWXV) downstream of an IPTG-inducible promoter (P_*tac*_-TWXV) and integrated this construct at an ectopic site. To assess the degree of overexpression, we generated a functional internally FLAG-tagged allele of PilX (*pilX*::FLAG) (**Fig. S3A**) to assess the relative expression level of minor pilins. Semi-quantitative western blots (**Fig. S4A-B**) of native *pilX*::FLAG vs *pilX*::FLAG P_*tac*_-TWX(::FLAG)V indicates that minor pilins are overexpressed in the latter when induced with 100 µM IPTG (**Fig. 2D**). Notably, while the *pilX*::FLAG allele is functional for natural transformation (**Fig. S3A**), it exhibits a slight reduction in the frequency of T4aP dynamics (**Fig. S3B-D**), which limits the sensitivity of these assays. As a result, all functional assays utilized strains with untagged *pilX*.

To determine whether minor pilin abundance regulates T4aP, we next assessed pilus dynamic activity in P_*tac*_-TWXV by epifluorescence timelapse microscopy. When uninduced, the frequency of dynamic activity was similar to the parent; however, when induced with 100 µM IPTG the frequency of dynamic activity for P_*tac*_-TWXV was dramatically increased (**Fig. 2A-C)**. Importantly, overexpression of the competence T4aP minor pilins (P_*tac*_-TWXV) did not markedly affect other properties of T4aP dynamics including pilus length, retraction rate, or the dwell time between extension and retraction (**Fig. S5A-C**). While a very modest, but significant, reduction was observed on T4aP extension rates (**Fig. S5D**), this difference is unlikely to explain the large increase in the frequency of dynamic activity observed. Together, these data strongly suggest that overexpression of the minor pilins increases the frequency of T4aP dynamic activity.

Thus far, we have only assessed the frequency of T4aP dynamics through direct observation by epifluorescence timelapse microscopy. Competence T4aP dynamic activity promotes the uptake of DNA across the outer membrane during natural transformation [16, 42]. Thus, if minor pilin overexpression increases the frequency of T4aP dynamic activity, we hypothesized that the rate of DNA uptake should increase in cells overexpressing minor pilins. To test this, we used a qPCR-based DNA uptake assay as a functional readout for T4aP dynamics. In brief, cells were incubated with a tDNA PCR product that contains a sequence that is unique from the host genome. At defined timepoints, aliquots of the reaction were removed and mixed with DNase I to degrade extracellular DNA and to prevent further tDNA uptake. Importantly, tDNA taken up across the outer membrane via competence T4aP dynamic activity is protected from degradation. DNase treated aliquots were then boiled (to inactivate the DNase and to release periplasmic tDNA via cell lysis) and used as template for qPCR reactions to detect the abundance of the tDNA product (see **Methods** for details). Altogether, this approach allowed us to kinetically monitor DNA uptake across the outer membrane (**Fig. 2E**). The line of best fit was determined by linear regression analysis to define the slope, which represents the rate of DNA uptake (**Fig. 2E-F**). Importantly, a strain incapable of assembling competence T4aP (Δ*pilA*) lacked any measurable uptake, which indicates that the tDNA uptake observed in this assay is dependent on T4aP dynamics. When we tested P_*tac*_-TWXV, we found that this strain exhibits a significant increase in the rate of DNA uptake compared to the parent (**Fig. 2E-F**). These results reinforce that overexpression of minor pilins dramatically increases the frequency of T4aP dynamics. Furthermore, these results demonstrate that the rate of DNA uptake serves as a functional correlate for the frequency of competence T4aP dynamics.

These results are consistent with a model wherein T4aP dynamics are constrained in the parent due to a limited supply of the minor pilin complexes needed to initiate T4aP assembly. Minor pilins and major pilins, however, are structurally similar and it is possible that these proteins compete with one another to gain access to the T4aP machine. Thus, a distinct possibility is that it is not the absolute abundance of minor pilins, but rather the stoichiometric ratio of minor:major pilins that regulates the frequency of T4aP dynamic activity. Overexpression of the minor pilins does not alter the absolute abundance of PilA, as determined by semi-quantitative western blots (**Fig. 2D, S4C-D**). Thus, overexpression of minor pilins via P_*tac*_-TWXV increases both the absolute abundance of minor pilins and the ratio of minor:major pilins, which prevents us from distinguishing between these two models.

### The stoichiometry of minor:major pilins regulates the frequency of T4aP dynamic activity

Next, we sought to increase the minor:major pilin ratio without increasing the absolute abundance of minor pilins to help discriminate between these two models. To do this, we sought to underexpress the major pilin while maintaining expression of minor pilins at their native level. If a low absolute abundance of minor pilins constrains T4aP dynamic activity in the parent, this perturbation should have a limited effect on the frequency of T4aP dynamics. However, if it is the ratio of minor:major pilins that regulates T4aP dynamics, then underexpression of the major pilin should increase the frequency of T4aP dynamics in a manner that phenotypically resembles overexpression of the minor pilins.

To test this hypothesis, we generated a strain with an in-frame deletion of the major pilin, *pilA*, and expressed *pilA in trans* via an IPTG-inducible P_*tac*_ promoter (pMMB::P_*tac*_-*pilA*). To assess whether this genetic background allows for titratable control of PilA expression without affecting expression of the minor pilins, we performed semi-quantitative western blots. These experiments showed that PilA levels were titratable in this background; with 50 µM IPTG inducing native levels of PilA, and lower concentrations of IPTG yielding underexpression of PilA (**S4C-E**). Importantly, minor pilin expression is unaffected in this background because PilX::FLAG levels were indistinguishable from the parent regardless of the IPTG concentration (**Fig. S4E**). These results indicate that Δ*pilA* pMMB::P_*tac*_-*pilA* will allow us to test our hypothesis because at concentrations of IPTG <50 µM the ratio of minor:major pilins is increased (**Fig. 3A**), while the absolute abundance of minor pilins remains unchanged (**Fig. S4E**).

**Fig. 3.**
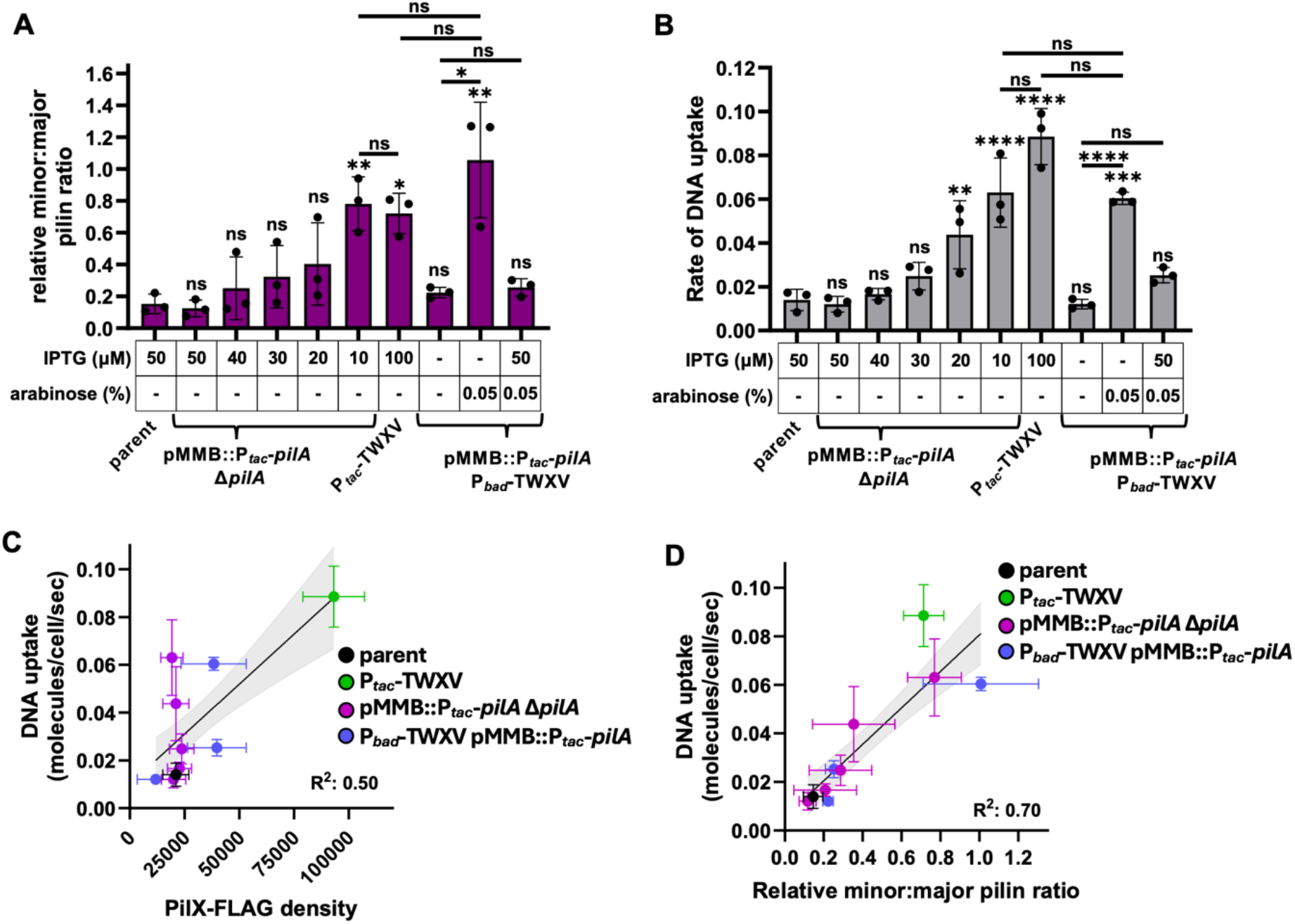
The stoichiometry of minor:major pilins, not the absolute abundance of minor pilins, regulates the frequency of T4aP dynamic activity. (**A**) The relative minor:major pilin ratios of the indicated strains grown in the presence of the indicated inducers were derived from semi-quantitative western blots (see **Fig. S4E** and **Dataset S1** for raw data). (**B**) Rates of DNA uptake for the indicated strains grown in the presence of the indicated inducers were derived from kinetic DNA uptake assays (see **Fig. S6** for raw data). The data for parent and P_tac_-TWXV in **A** and **B** are identical to the those presented in **Fig. 2** and are included here for ease of comparison. Statistical comparisons in **A** and **B** were made by one-way ANOVA with Tukey’s multiple comparison test of the log-transformed data. NS = no significance, * = p < 0.05, ** = p < 0.01, *** = p < 0.001, **** = p < 0.0001. Statistical identifiers directly above bars represent comparisons to the parent. For all of the data collected in the manuscript, linear regression analysis was performed to separately assess the correlation between the rate of DNA uptake and (**C**) PilX::FLAG density and (**D**) the minor:major pilin ratio. Lines of best fit with 95% confidence bands (gray fill) and R^2^ values are shown. All data in **C** and **D** are identical to those presented in **A, B**, and **Fig. S4E**, and are from at least three independent biological replicates and shown as the mean ± SD.

When PilA is expressed at parent levels in this background (50 µM IPTG), the frequency of T4aP dynamic activity is indistinguishable from the parent as assessed by timelapse epifluorescence microscopy (**Fig. S6A-B**) and the rate of DNA uptake (**Fig. 3B, S6C**). Underexpression of PilA in this background (10 µM - 40 µM IPTG), however, results in a progressive increase in observable T4aP dynamics (**Fig. S6A-B**) and DNA uptake (**Fig. 3B, S6C**). Importantly, the increase in T4aP dynamics observed by timelapse epifluorescence microscopy directly correlates with the rate of DNA uptake (**Fig. S6E**), which further reinforces that the rate of DNA uptake serves as a functional correlate for the frequency of competence T4aP dynamics. When PilA is underexpressed using 10 µM IPTG, the relative minor:major pilin ratio is similar to when minor pilins are overexpressed (**Fig. 3A**), and the frequency of T4aP dynamics is statistically indistinguishable between these conditions (**Fig. 3B**). Importantly, underexpression of PilA does not affect pilus length, retraction rate, or the dwell time between extension/retraction; and only modestly decreases the extension rate, which is consistent with what is observed when minor pilins are overexpressed (**Fig. S5A-D**). Also, the increase in the frequency of T4aP dynamics observed when the major pilin is underexpressed is dependent on minor pilins (**Fig. S5E**).

Thus far, our experiments have shown that both 1) overexpressing the minor pilins and 2) underexpressing the major pilin results in a dramatic increase in dynamic activity, supporting the hypothesis that the minor:major pilin ratio, not the absolute abundance of minor pilins, regulates the frequency of T4aP dynamic activity. To further test this hypothesis, we sought to increase minor pilin abundance while maintaining the minor:major pilin ratio at parent levels. If the minor:major pilin ratio regulates the frequency of T4aP dynamics, we would hypothesize that this should result in T4aP dynamics that are similar to the parent (despite the increase in abundance of minor pilins). To that end, we generated a strain where minor pilin overexpression is regulated by an arabinose inducible promoter (P_*bad*_-TWXV) and major pilin overexpression is regulated by IPTG-inducible pMMB::P_*tac*_-*pilA* in a background where the native minor pilin locus and *pilA* are intact (which allows for inducible overexpression of both the minor and major pilins). When pMMB::P_*tac*_-*pilA* P_*bad*_-TWXV is uninduced, it is indistinguishable from the parent for both the minor:major pilin ratio (**Fig. 3A, S4E**) and the frequency of T4aP dynamics (**Fig. 3B**). When pMMB::P_*tac*_-*pilA* P_*bad*_-TWXV is induced with only arabinose, only the minor pilins are overexpressed (**Fig. S4E**), which yields an increase in the minor:major pilin ratio (**Fig. 3A**) and a corresponding increase in the frequency of T4aP dynamics akin to what was previously observed for P_*tac*_-TWXV (**Fig. 3B, S6D**). When pMMB::P_*tac*_-*pilA* P_*bad*_-TWXV is induced with both arabinose and IPTG, however, both the minor and the major pilins are similarly overexpressed (**Fig. S4E**), which yields a minor:major pilin ratio that is similar to the parent (**Fig. 3A**) and a frequency of T4aP dynamics that was also indistinguishable from the parent (**Fig. 3B, S6D**). These results further support the hypothesis that it is the minor:major pilin ratio, not the absolute abundance of minor pilins, that regulates the frequency of T4aP dynamic activity.

To formalize this comparison, we sought to quantitatively assess whether the frequency of T4aP dynamics is more strongly correlated with 1) minor pilin abundance (**Fig. 3C**) or 2) the minor:major pilin ratio (**Fig. 3D**) using all of the data collected. This analysis revealed a poor correlation between the frequency of T4aP dynamic activity and minor pilin abundance (**Fig. 3C**); and a comparably stronger correlation between the frequency of T4aP dynamic activity and the minor:major pilin ratio (**Fig. 3D**). All together, these data strongly support a model wherein the minor:major pilin ratio, not the absolute abundance of minor pilins, regulates the frequency of T4aP dynamic activity.

## DISCUSSION

This study sheds light on a mechanism used to regulate dynamic activity in T4aP systems. Specifically, we show that the stoichiometric ratio of minor:major pilins, not their absolute abundance, regulates the frequency of competence T4aP dynamics in *V. cholerae*. The molecular mechanism underlying this regulation is unclear, however we suggest two possibilities. The core minor pilins likely interact with one another to form initiation complexes in the membrane, thus, one possibility is that a high minor:major pilin ratio simply increases the likelihood that initiation complexes will form; while a low minor:major pilin ratio disrupts assembly due to the inappropriate incorporation of the major pilin into nascent initiation complexes. Following the formation of initiation complexes, the minor pilins must interact with the T4aP machine to prime filament extension. So, a second possibility is that a higher minor:major pilin ratio allows initiation complexes to more successfully compete with the major pilin for access to the T4aP machine, thus, priming more T4aP machines for extension. Distinguishing between these two possibilities is experimentally challenging and will likely require in-depth structural and/or biochemical analysis *in situ* to determine how the minor:major pilin ratio impacts assembly of the minor pilin complex and the subsequent interaction between this complex and the T4aP machine. T4aP are only one subset of the broader type IV filament family [3]. While minor pilins likely promote assembly of other T4P (including the T4bP, T4cP, and T4dP) important differences between the structure / number of minor pilins, T4P machine components, and retraction mechanisms among T4P make it unclear whether the minor:major pilin ratio will have a similar regulatory role in these systems.

Our findings add to a growing list of factors that have been shown to influence the frequency of T4aP dynamics in diverse bacterial species. In some T4aP systems, the activity of the extension motor ATPase is allosterically regulated, either directly or indirectly, via the secondary messenger cyclic-di-GMP [49-53]. The extension and retraction motor ATPases may also antagonistically compete for the T4aP machine in *P. aeruginosa* [36], suggesting that the ratio of motor ATPases can alter the frequency of dynamic activity. Also, T4aP dynamics are spatially and temporally regulated in many systems, especially for twitching motility. This includes regulation of twitching motility by a chemosensory system in *P. aeruginosa* [32, 54], phototaxis in cyanobacteria [55], and the control of social motility in *M. xanthus* [56]. Here, we show that the minor:major pilin ratio can also influence the frequency of T4aP dynamics. The use of four core minor pilins to initiate pilus assembly is a universally conserved feature of T4aP in diderms [4, 6]. Thus, it is tempting to speculate that the minor:major pilin ratio affects the dynamic activity of T4aP in other species. Also, many species encode the PilS/PilR two-component system that transcriptionally regulates the expression of the major pilin based on the pilin load of the inner membrane [57-61]. This could help stabilize minor:major pilin ratios for consistent dynamic activity; or perhaps more speculatively, could provide a mechanism for altering minor:major pilin ratios to tune dynamic activity based on environmental cues. Notably, *V. cholerae* lacks PilS/PilR; and our results demonstrate that expression of the competence T4aP minor and major pilins is uncoupled, which allowed us to easily manipulate the relative minor:major pilin ratio in this study.

Our findings suggest that the minor:major pilin ratio regulates the frequency of T4aP dynamics. Other properties of T4aP dynamics include the rate of extension/retraction, the length of pili, and the dwell time between extension/retraction. Prior work has shown that extension/retraction rates are defined, in part, by the ATP hydrolysis rates of the motor ATPases that power extension/retraction, respectively [30, 33, 39, 62]. We show here that the minor:major pilin ratio also has a slight, but significant, negative effect on extension rates; however, a mechanistic understanding for this observation remains lacking. A preprint posted while this publication was under review suggests that major pilin abundance regulates pilus length in *P. aeruginosa* [63], however, our results suggest that this is not a conserved property of *V. cholerae* competence T4aP. Thus, T4aP length may be governed by distinct mechanisms in different systems. Also, the factors that define the switch between extension and retraction remain poorly understood in most T4aP systems, which will be the focus of future work.

Our results highlight that the frequency of competence T4aP dynamics are constrained in *V. cholerae*. But why would bacteria want to constrain the dynamic activity of their T4aP? Our results show that increasing T4aP dynamics can increase the rate of DNA uptake for the competence T4aP in *V. cholerae*. Thus, increased T4aP dynamics can clearly benefit cells by enhancing their capacity for horizontal gene transfer by natural transformation. On the other hand, increased T4aP dynamics may incur a fitness cost. First, T4aP dynamics are likely energetically costly. Structural modeling of the motor ATPases that drive extension/retraction indicate that the addition/removal of a single pilin subunit likely requires hydrolysis of 2 ATP [64, 65]. Because T4aP filaments are often composed of thousands of pilin subunits, increased dynamics could drain cellular resources. Also, T4aP are common phage receptors. So increased T4aP dynamics can increase the vulnerability to phage infection. One way bacteria can mitigate this vulnerability is by reducing the frequency of T4aP dynamics [53]. Also, recent studies highlight that prophages decrease T4aP dynamics as a mechanism of superinfection exclusion [66, 67], which further suggests that constraining T4aP dynamics can promote phage resistance. Finally, there may be environmental conditions where T4aP-dependent activities are disadvantageous. For example, retraction of MSHA T4aP and cessation of dynamics promotes escape from environmental stressors [41], and enhances colonization of mammalian hosts [68] in *V. cholerae*. We find here that T4aP dynamics are influenced, in part, by the ratio of minor:major pilins. This simple regulatory mechanism may allow bacteria to rapidly optimize T4aP dynamics to balance the associated costs and benefits.

## METHODS

### Bacterial strains and culture conditions

All *V. cholerae* strains were derived from *V. cholerae* E7946 [69]. Cells were routinely grown in LB Miller broth rolling at 30 °C or on LB miller agar at 30 °C supplemented with trimethoprim (10 µg/mL), spectinomycin (200 µg/mL), kanamycin (50 µg/mL), tetracycline (0.5 µg/mL), erythromycin (10 µg/mL), chloramphenicol (1 µg/mL), carbenicillin (100 µg/mL) or zeocin (100 µg/mL) as appropriate.

In order to isolate and study dynamic activity of the competence T4aP in *V. cholerae*, the parent strain used throughout this study had several critical mutations: 1) Expression of competence T4aP naturally requires chitin oligosaccharides and quorum sensing as inducing cues. To reliably study T4aP dynamics, we bypassed these native signals by overexpressing the master regulator of competence, TfoX, via a constitutively active promoter, P_const2_, and deleted *luxO* to genetically lock cells in a high density state as previously described [16]. 2) To detect active T4aP using fluorescence microscopy, all strains harbored a *pilA*^*S67C*^ mutation which allows for pilus labeling with fluorescent maleimide dyes [16, 40]. 3) *V. cholerae* strains naturally encode two plasmid defense systems, *ddmABC* and *ddmDE*, that can deplete cells of plasmids used for ectopic complementation [70]. To use pMMB::P_tac_-*pilA* in the absence of antibiotics, these two plasmid defense systems were deleted.

### Construction of mutant strains

All strains were generated by natural transformation using mutant constructs assembled by splicing-by-overlap extension (SOE) exactly as previously described [71, 72] and confirmed by PCR and sequencing. For a detailed list of strains used for this study, see **Table S1**. For a complete list of primers used to generate mutant constructs, see **Table S2**.

The pMMB::P_tac_-*pilA* plasmid was assembled by SOE PCR and cloned into chemically competent *Escherichia coli* TG1. This plasmid was then miniprepped and introduced into *V. cholerae* via natural transformation.

### Direct visualization of competence T4aP by epifluorescence microscopy

To prepare cells for imaging, cultures were grown at 30 °C rolling in LB supplemented with 10-100 µM IPTG, 20 mM MgCl_2_, 10 mM CaCl_2_ to late log. Then, ∼10^8^ cells were harvested and washed in instant ocean medium (IO; 7g/L, Aquarium Systems) to remove residual LB. To label competence T4aP, washed cells were incubated with 25 µg/mL AlexaFluor 488-maleimide (AF488-mal) dye (ThermoFisher) for 20-30 mins at room temperature. To remove unbound dye, cells were washed three times in IO. Then, cells were placed under a 0.4% gelzan pad for imaging. Phase contrast and widefield fluorescence images were collected using a Nikon Ti-2 microscope using a Plan Apo × 60 objective, a FITC filter cube, a Hamamatsu ORCA Flash 4.0 camera and Nikon NIS Elements imaging software. For timelapse imaging, samples were incubated at 30 °C using an adjustable objective warmer (Bioptechs Inc.).

### Microscopy image analysis

To assess surface piliation in Δ*pilT* strains, static images were taken of AF488-mal labeled cells. Fiji imaging software [73] with the microbeJ plugin [74] was used to quantify the number of cells in a field of view. Then, the fields of view were visually scanned to quantify the number of cells with rare piliation events (in the case of the minor pilin deletion mutants) or rare non-piliated cells (in the case of the Δ*pilT* parent or minor pilin complemented strains).

To quantify the frequency of dynamic activity, timelapse microscopy was performed by imaging AF488-mal labeled cells every 3 sec for 2 mins. Samples were incubated at 30 °C for the duration of the timelapse. Timelapse images were analyzed using NIS elements software. For each replicate, 60 cells were randomly selected using the phase contrast channel. For each cell, the fluorescence channel was then manually analyzed to determine the number of extension and/or retraction events observed. Every dynamic pilus observed within the 2 min timelapse was scored as a distinct dynamic event. We have found empirically that pili < 0.15 µm cannot be resolved via this imaging approach.

The maximum length of dynamic competence T4aP was measured using the same timelapse images used to quantify dynamic activity. NIS elements software was used and cells in a field of view were analyzed one-by-one for piliation events during the 2 min timelapse. When a pilus was observed, its maximum length was measured using the multi-point tool to account for the flexibility of T4aP.

To measure extension rates, retraction rates, and dwell times, timelapse microscopy was performed by imaging AF488-mal labeled cells every 1 sec for 1 min. Cells were visually assessed to identify pili that completely extended and retracted during the course of the timelapse. The frame and timestamp at the beginning and end of the extension/retraction event was recorded. Also, the maximum pilus length for each dynamic event was measured. Then, the length of the pilus (in µm) was divided by the elapsed time for extension and retraction (in sec) to define the extension rate and retraction rate (µm/s), respectively. The dwell time was defined as the time (in sec) elapsed between when a pilus stopped extending and when retraction began.

### Semi-quantitative western blot analysis

Strains with P_*tac*_-TWXV or pMMB::P_*tac*_-*pilA* only were grown as described above for microscopy. Strains with P_*bad*_-TWXV and pMMB::P_*tac*_-*pilA* were grown in the same conditions, but with 50 µM IPTG and/or 0.05% arabinose, as indicated. For PilA, ∼5x10^9^ cells were centrifuged and resuspended in 50 µL IO. This was then mixed 1:1 with 2x SDS-PAGE sample buffer (200 mM Tris pH 6.8, 25% glycerol, 4.1% SDS, 0.02% Bromophenol Blue, 5% β-mercaptoethanol) and boiled for 10 mins. Samples were diluted into a Δ*pilA* lysate (*i*.*e*., a lysate lacking any FLAG-tagged proteins or PilA) so that the signal was within the linear range of the assay (see **Fig. S4**), and then electrophoretically separated on a 15% SDS-PAGE gel. Samples were diluted into a Δ*pilA* lysate (as opposed to loading buffer) to ensure that differences in protein abundance between samples did not impact the western blotting procedure (*i*.*e*., how samples ran on the gel, transfer to the membrane, etc.).

For PilX::FLAG, ∼5x10^10^ cells were centrifuged and resuspended in 50 µL IO. The solution was then mixed with lysis buffer [1x FastBreak lysis buffer (Promega), 3.8 µg/mL lysozyme (GoldBio), and 25 U Benzonase (Sigma Aldrich)] and incubated at room temperature for 10 mins. Then, each lysis reaction was mixed 1:1 with 2x SDS-PAGE sample buffer. Samples were diluted into a lysate of the parent (no FLAG) so that the signal was within the linear range of the assay (see **Fig. S4**). Samples were then electrophoretically separated on a 15% SDS-PAGE gel.

Proteins were then electrophoretically transferred to PVDF membranes. To detect PilA, blots were stained with polyclonal α-PilA primary antibodies [38]. To detect PilX::FLAG, blots were stained with monoclonal α-FLAG primary antibodies (Sigma). Blots were then incubated with HRP-conjugated secondary antibodies, developed using enhanced chemiluminescence (ECL) western blotting substrate (Pierce), and imaged on a ProteinSimple Fluorechem R instrument.

Protein densities were quantified using Fiji imaging software. Serial dilutions of PilX::FLAG and PilA lysates were performed to identify the linear range of detection for these western blots (see **Fig. S4**). Experimental samples were diluted so that their signal density was within the linear range (see **Fig. S4**), and densities were then multiplied by the dilution factor to obtain the signal density of the original sample.

### qPCR-based DNA uptake assays

Strains with P_*tac*_-TWXV or P_*tac*_-*pilA* only were grown as described above for microscopy. Strains with P_*bad*_-TWXV and P_*tac*_-*pilA* were grown in the same conditions, but with either 50 µM IPTG, 0.05% arabinose, or a combination of IPTG and arabinose. Each DNA uptake reaction consisted of IO, 0.1 mg/mL recombinant BSA (NEB), 2 ng/µL transforming DNA (tDNA; a 300 bp Δ*VC1102* PCR product), and ∼1x10^7^ cells. Each reaction was incubated at 30°C, and aliquots were taken at 30, 60, 120, 240, and 480 seconds after the addition of tDNA. At each timepoint, the aliquot was immediately added to a DNase solution consisting of IO, 0.1 mg/mL recombinant BSA, 1x DNase buffer (NEB), and 6 units of DNase I (NEB) and mixed by pipetting. After allowing the DNase reaction to proceed at room temperature for 2 mins, samples were boiled and then placed on ice.

Next, each sample was used as template for qPCR on a StepOnePlus instrument (Applied Biosystems). To determine the quantity of tDNA taken up, qPCR was performed with a primer pair that would uniquely detect the tDNA. And to normalize samples, qPCR was also performed for *rpoB* to quantify the number of cells in each aliquot. The tDNA molecules per cell was determined by dividing the quantity of Δ*VC1102* by the quantity of *rpoB*. Then, linear regression analysis was performed to determine the rate of DNA uptake for each strain.

### Structural modeling of minor pilins

Proteins were modeled using the AlphaFold-multimer algorithm [75] in ColabFold [76] on Indiana University’s high-performance computing (HPC) system. Colabfold runs were configured for 10 recycles with an early stop tolerance of 0.4. Models were analyzed and visualized using UCSF ChimeraX [77].

### Statistics

Statistical differences were assessed using GraphPad Prism software. The statistical tests used are indicated in the figure legends. Descriptive statistics for all samples, and a comprehensive list of statistical comparisons can be found in **Dataset S1**.

## ACKNOWLEDGEMENTS

This work was supported by Grant R35GM128674 from the NIH to A.B.D.

---

**Fig. S1.**
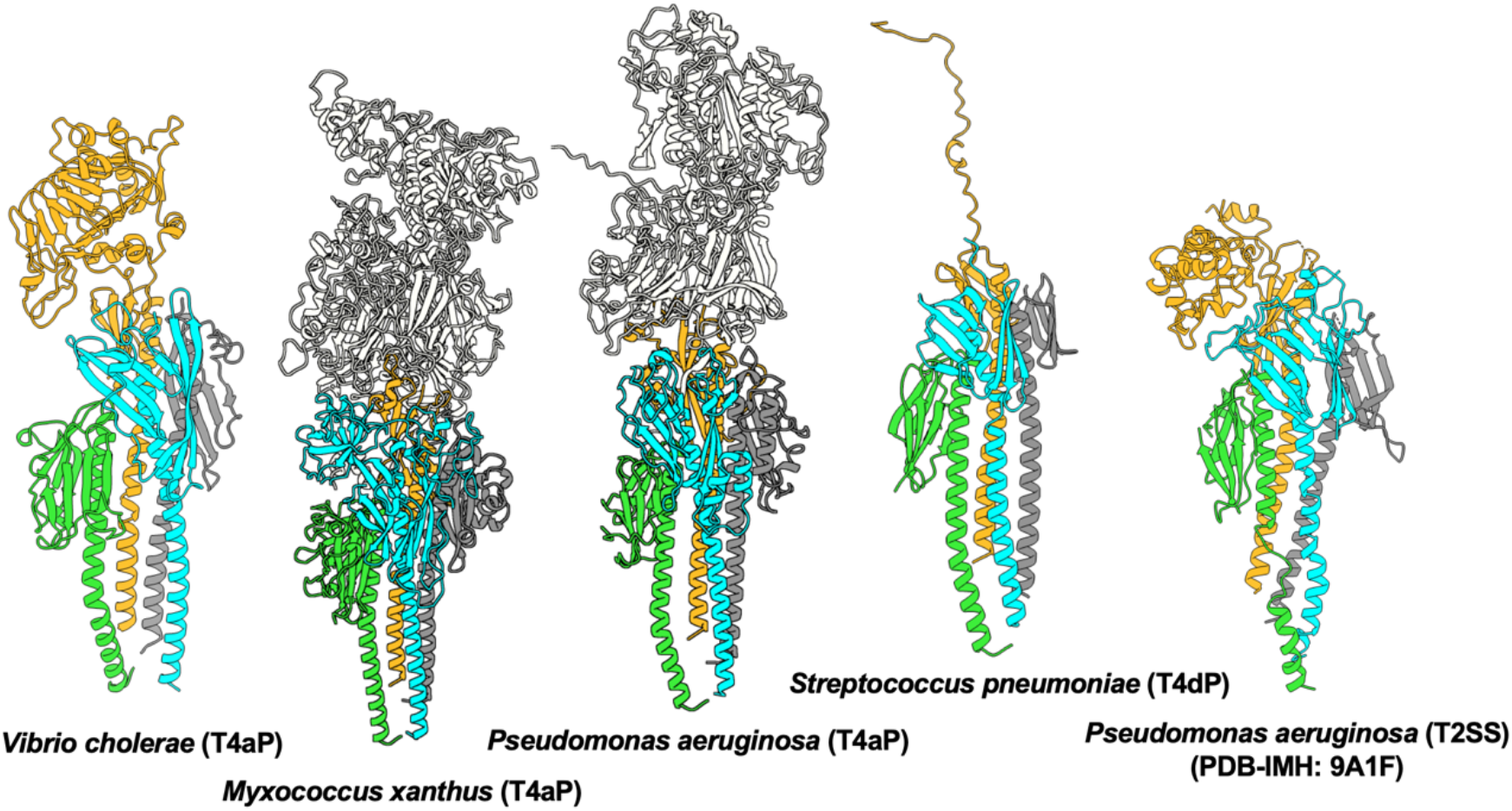
The core minor pilins from diderm T4aP, T4dP, and T2SS share a conserved structural arrangement. AlphaFold-multimer models of the core minor pilins from *V. cholerae* (T4aP), *M. xanthus* (T4aP; Treuner-Lange, Chang et al. 2020), *P. aeruginosa* (T4aP ; Nguyen, Sugiman-Marangos et al. 2015), *S. pneumoniae* (T4dP; Christman and Dalia 2025), and *P. aeruginosa* (T2SS; Escobar et al., 2021). Pilins are colored based on their structure and position within the predicted complex. Green = FimU/T, Cyan = PilW, Grey = PilV, Yellow = PilX. For both *M. xanthus* and *P. aeruginosa* T4aP, PilY1 is included in white.

**Fig. S2.**
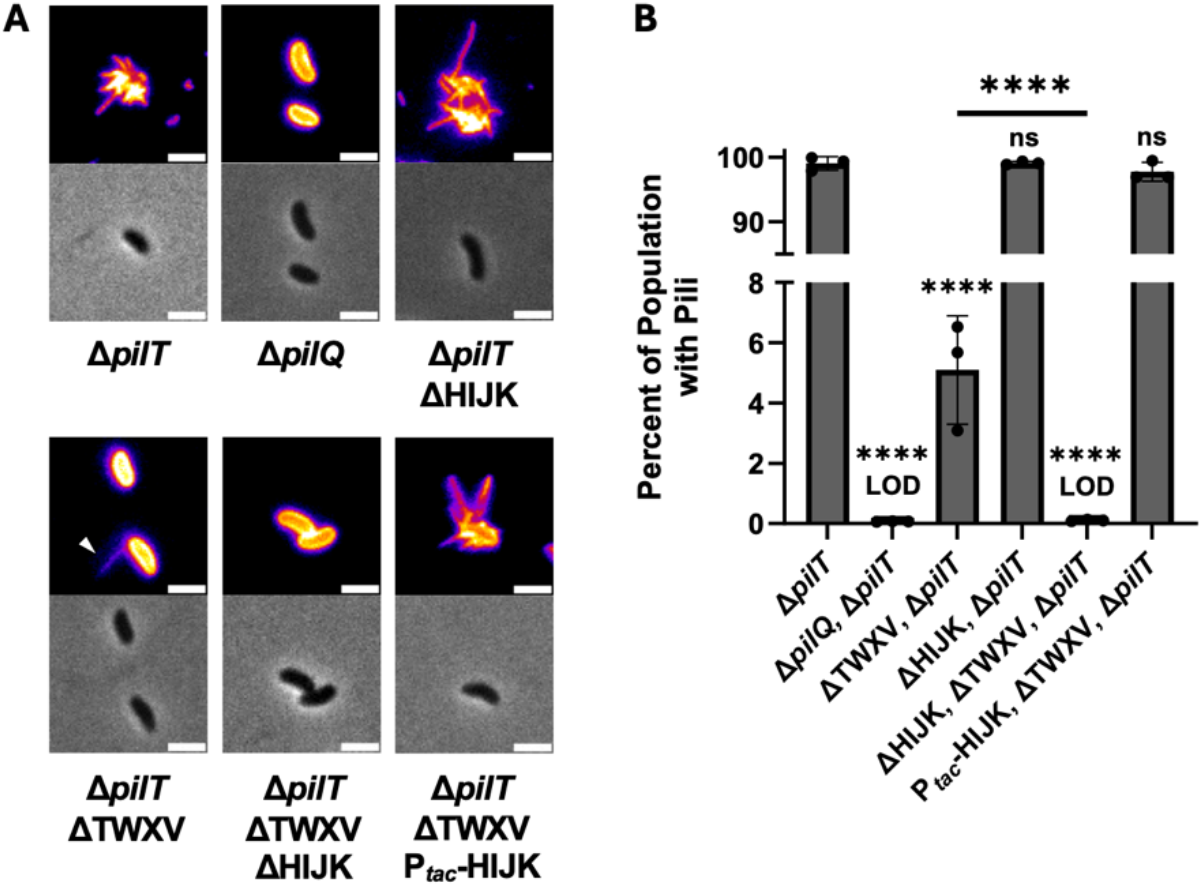
T2SS minor pilins can promote assembly of the competence T4aP in *V. cholerae*. (**A**) Representative images of the indicated strains stained with AF488-mal. All strains were induced with 100 µM IPTG. Images are false colored with the “Fire” LUT in Fiji to make rare pili easier to see (white arrows). Scale bar, 2 µm. (**B**) Quantification of surface piliation in the indicated strains stained with AF488-mal, as depicted in **A**. For these assays, the parent strain is Δ*vesC* Δ*epsM*. The Δ*vesC* mutation suppresses the lethality typically associated with loss of T2SS activity in *V. cholerae*, and the Δ*epsM* mutation disrupts an essential T2SS machine component to ensure that all strains lack T2SS activity. Data in **B** are from 3 independent biological replicates and shown as the mean ± SD. The number of cells quantified varied for each strain and replicate. The total number of cells analyzed for each strain: Δ*pilT* = 3067, Δ*pilT* Δ*pilQ* = 4523, Δ*pilT* ΔHIJK = 3123, Δ*pilT* ΔTWXV = 4273, Δ*pilT* ΔTWXV ΔHIJK = 3131, Δ*pilT* ΔTWXV P_tac_-HIJK = 2647. Statistical comparisons were made by one-way ANOVA with Tukey’s multiple comparison test of the log-transformed data. NS, no significance; **** = *p* < 0.0001. LOD, limit of detection. Statistical identifiers directly above bars represent comparisons to Δ*pilT*.

**Fig. S3.**
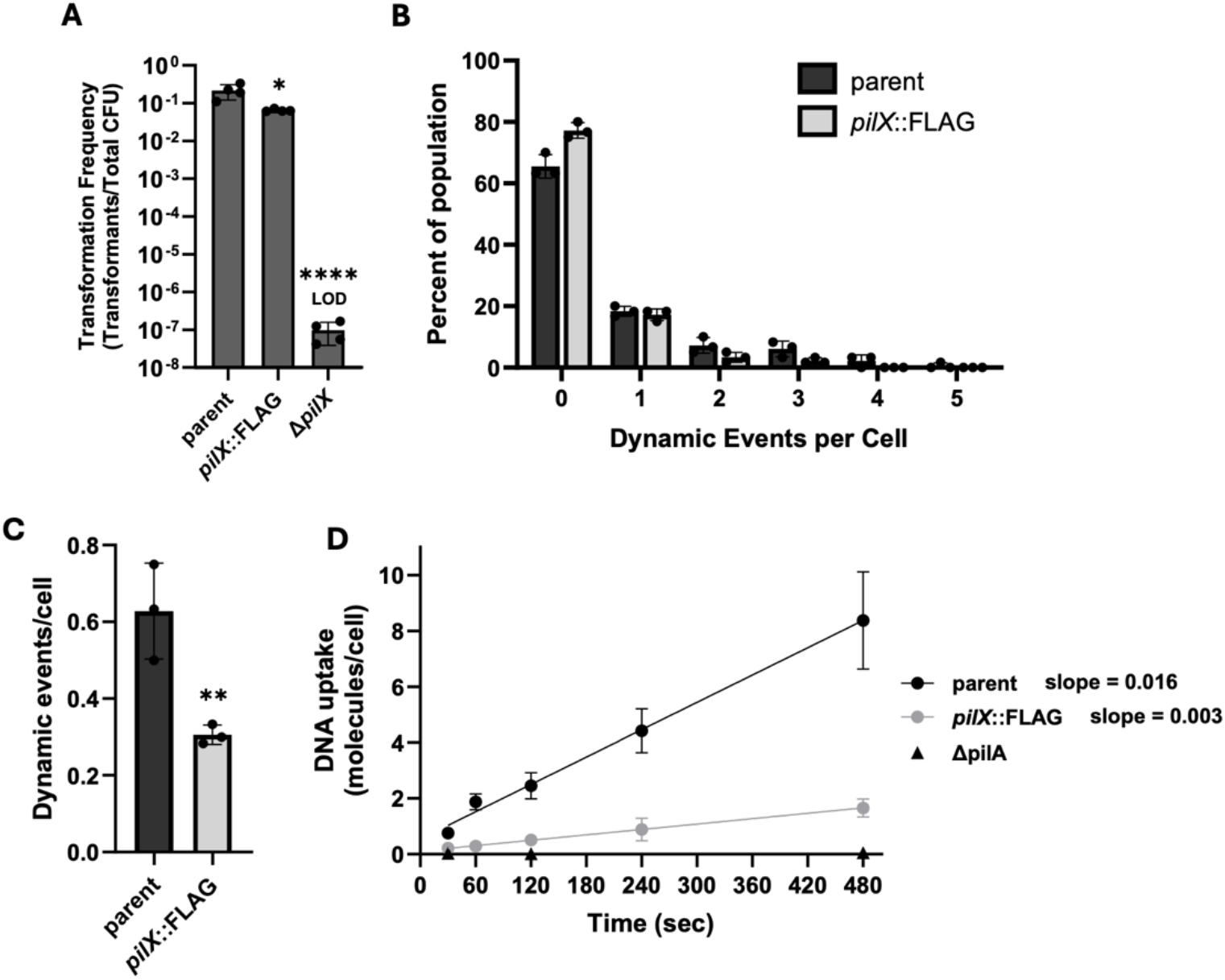
PilX::FLAG is functional but exhibits slightly reduced dynamic activity. (**A**) Transformation assays of the indicated strains. (**B**-**C**) The frequency of T4aP dynamics of the indicated strains was quantified from timelapse imaging of AF488-mal labeled cells and the (**B**) distribution and (**C**) average T4aP dynamics per cell are plotted. Parent data shown here are identical to data shown in **Fig. 2B and 2C** and are included here for ease of comparison. (**D**) DNA uptake into the periplasm of the indicated strains was kinetically monitored by qPCR. Parent and Δ*pilA* are identical to the data presented in **Fig. 2E** and are included here for ease of comparison. Lines of best fit were determined by linear regression. The slope of each line is shown, which represents the rate of DNA uptake. Data is from at least 3 independent biological replicates and shown as the mean ± SD. Statistical comparisons were made in **A** by one-way ANOVA with Tukey’s multiple comparison test of the log-transformed data, and in **C** by unpaired Student’s t-test of the log-transformed data. * = *p* < 0.05, ** = *p* < 0.01, **** = *p* < 0.001. LOD = limit of detection. Statistical identifiers directly above bars represent comparisons to the parent.

**Fig. S4.**
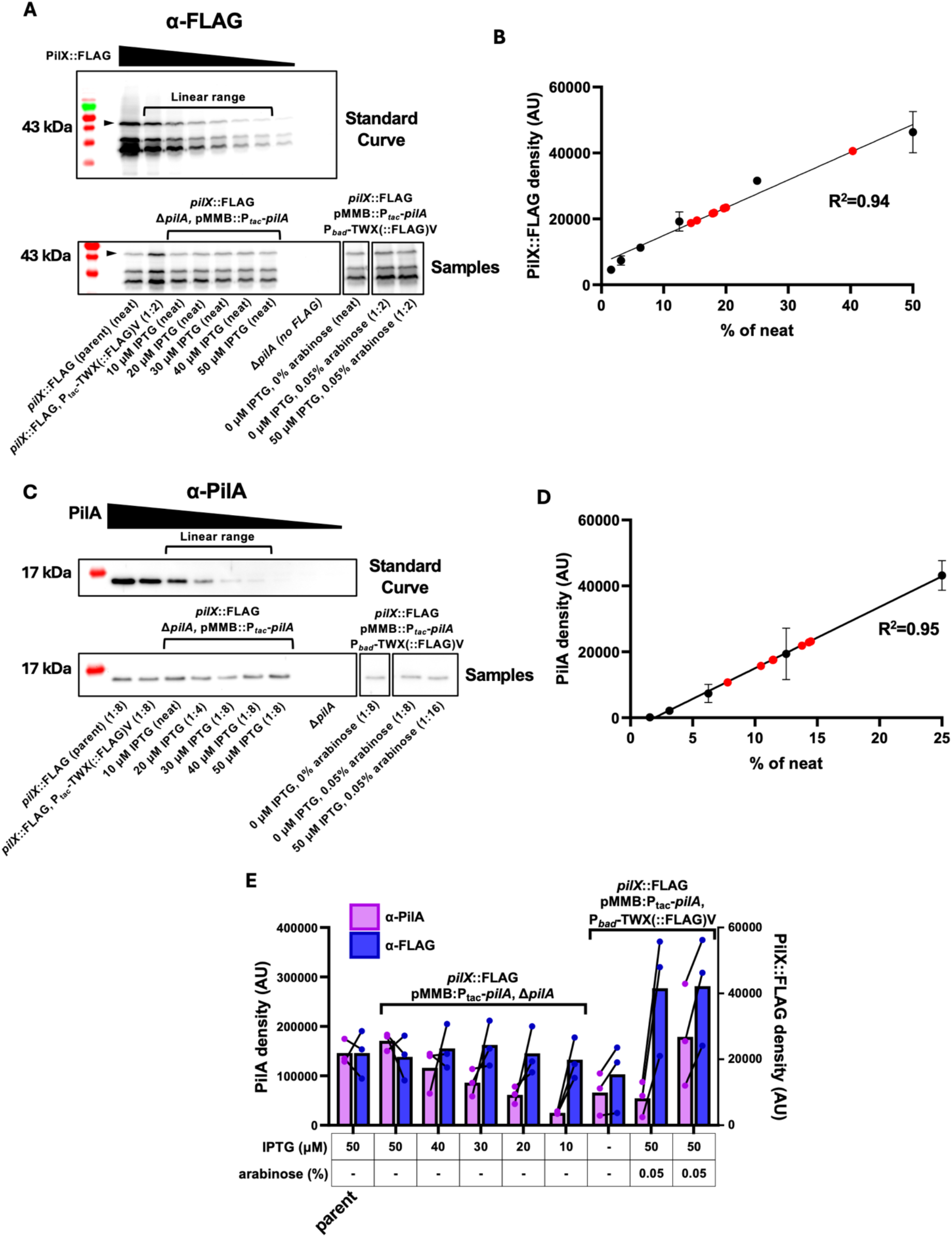
Assessing minor and major pilin levels via semi-quantitative western blots. (**A**) Representative semi-quantitative western blots for detecting PilX::FLAG using α-FLAG primary antibodies. A serial 2-fold dilution series of PilX::FLAG (*i*.*e*., dilutions of a *pilX*::FLAG P_*tac*_-TWX(::FLAG)V induced with 100 µM IPTG lysate) (top) was used to generate a standard curve to define the linear range of the assay. Experimental samples (bottom) from the indicated strains grown with the indicated concentration of inducer were diluted (dilution factor noted in parentheses) and blotted so that the signal would be within the linear range of the assay. The band that corresponds to full length PilX::FLAG (45 kDa) is demarcated with a black arrow, and the size of the closest ladder band is noted to the left of the blot. (**B**) Blots, as shown in **A**, were analyzed by densitometry and plotted. Black datapoints correspond to the standard curve (*n* = 3), and only data points within the linear range are shown. Linear regressions show the line of best fit for the standard curve and the R^2^ is shown on the plot. Red datapoints correspond to the densities of a subset of experimental samples (as depicted in **A)** to highlight that these data fall within the linear range of the assay. (**C**) Representative semi-quantitative western blots for detecting PilA via α-PilA antibodies. A 2-fold dilution series of PilA (*i*.*e*., dilutions of a Δ*pilA* pMMB::P_*tac*_-*pilA* lysate) (top) was used to generate a standard curve to define the linear range of the assay. Experimental samples (bottom) from the indicated strains grown with the indicated concentration of inducer were diluted (dilution factor noted in parentheses) and blotted so that the signal would be within the linear range of the assay. (**D**) Blots, as shown in **C**, were analyzed by densitometry and plotted. Black datapoints correspond to the standard curve (*n* = 3), and only data points within the linear range are shown. Linear regressions show the line of best fit for the standard curve and the R^2^ is shown on the plot. Red datapoints correspond to the densities of a subset of experimental samples (as depicted in **C)** to highlight that these data fall within the linear range of the assay. PilA is 15 kDa, and the size of the closest ladder band is demarcated to the left of the blot. (**E**) Quantification of α-PilA and α-FLAG from semi-quantitative westerns of the indicated strains grown in the indicated inducers (*n* = 3). Paired data points represent samples from the same biological replicate. Parent data is identical to that in **Fig. 2D** and is included here for ease of comparison.

**Fig. S5.**
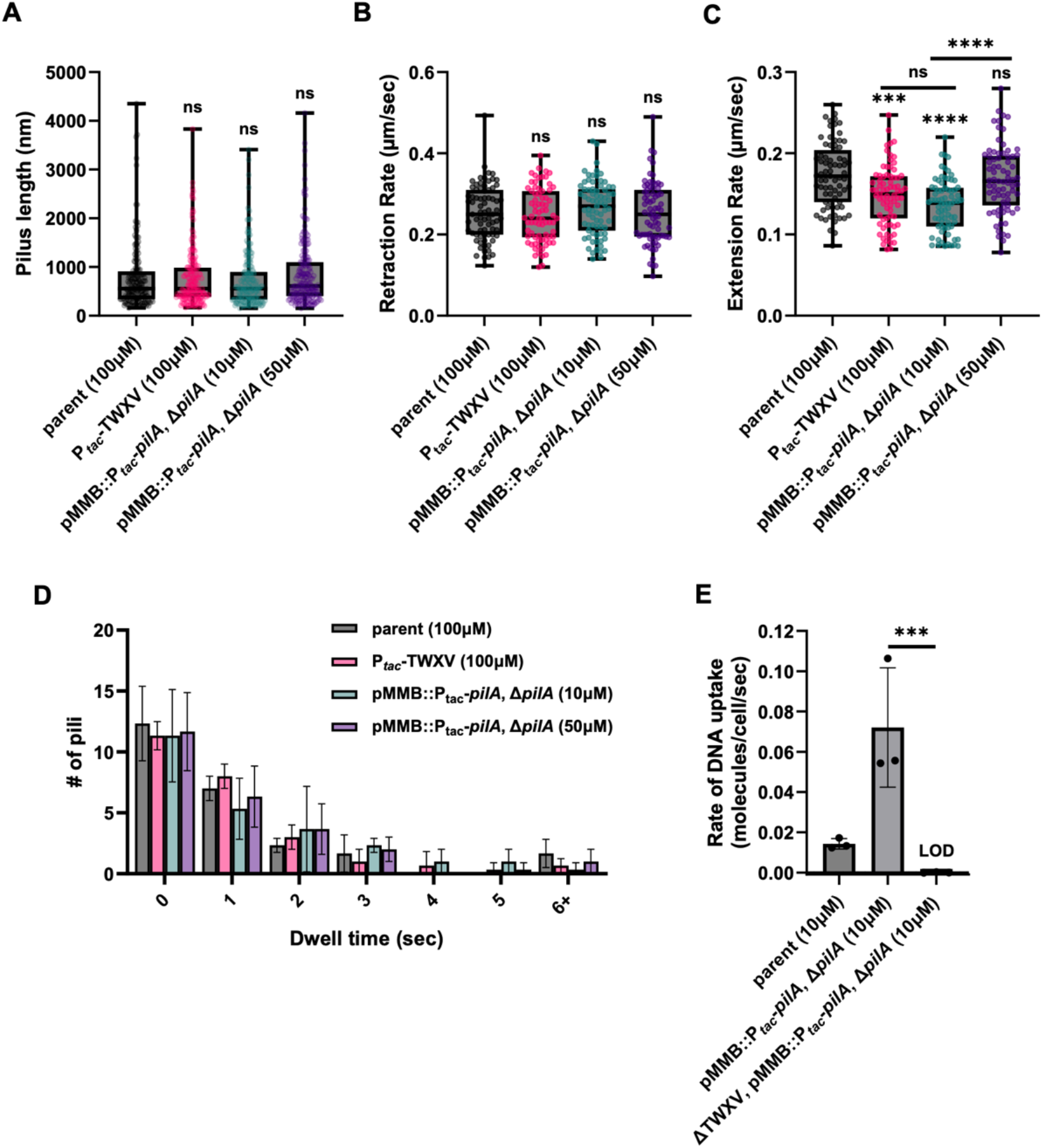
The minor:major pilin ratio does not affect other properties of T4aP dynamics. Timelapse imaging was performed on AF488-mal labeled cells of the indicated strains grown in the indicated concentration of IPTG and images were analyzed to determine the following properties of T4aP dynamics. (**A**) Maximum pilus lengths. (**B**) The rate of pilus retraction. (**C**) The rate of pilus extension. (**D**) The dwell time for pili. Dwell time is defined as the amount of time before an extended pilus begins to retract. (**E**) Rates of DNA uptake for the indicated strains derived from kinetic DNA uptake assays. All data are from 3 independent biological replicates and shown as the mean ± SD. Statistical comparisons were made by one-way ANOVA with Tukey’s multiple comparison test of the log-transformed data. ns = no significance, *** = *p* < 0.001, **** = *p* < 0.0001. Statistical identifiers directly above bars represent comparisons to the parent.

**Fig. S6.**
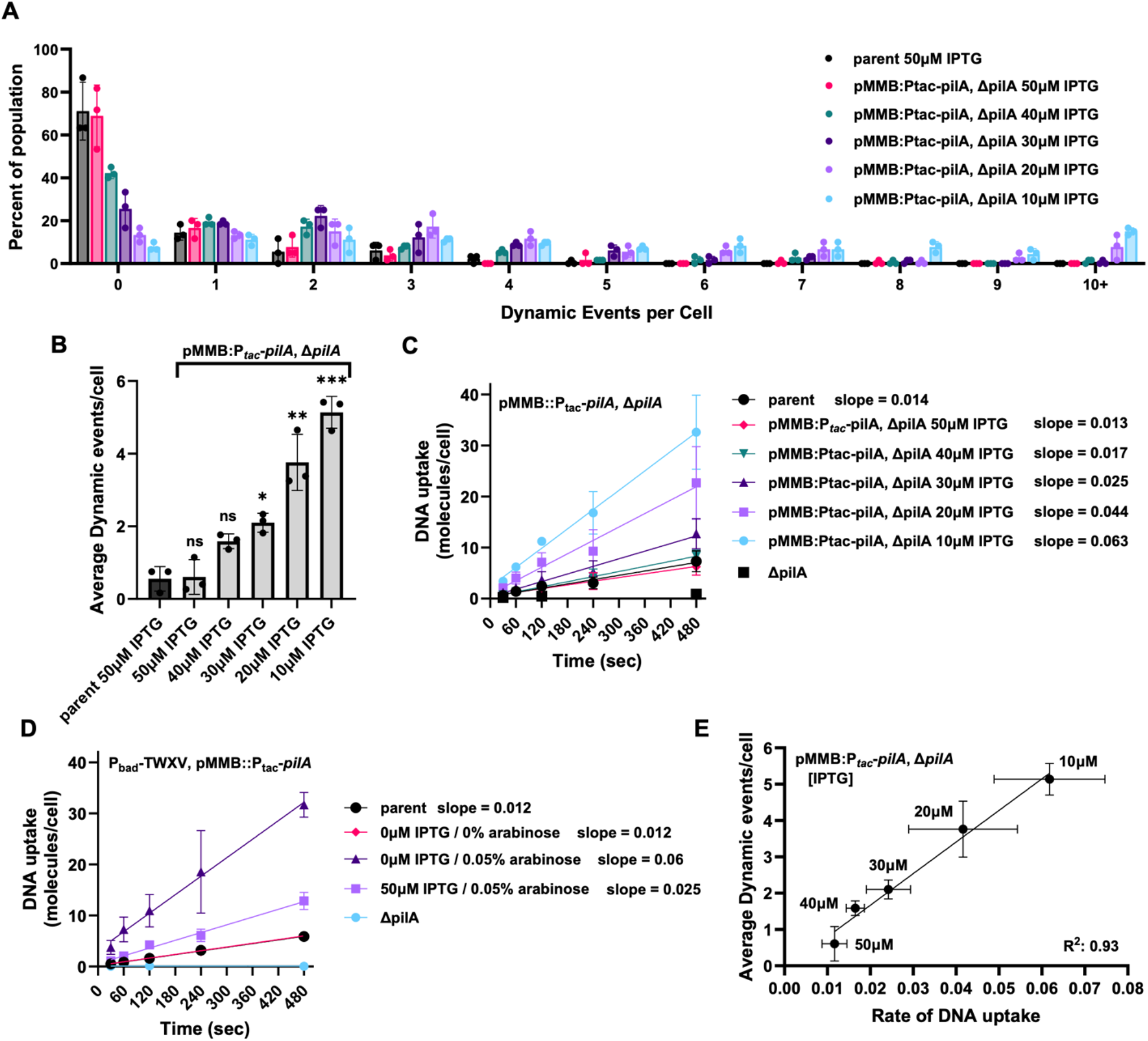
Raw data for quantification of the frequency of T4aP dynamic activity when the minor:major pilin ratio is experimentally modulated. (**A-B**) Quantification of the frequency of T4aP dynamics from timelapse imaging of the indicated strains stained with AF488-mal with (**A**) distribution and (**B**) average dynamics per cell shown. (**C**-**D**) DNA uptake into the periplasm of the indicated strains grown in the indicated inducers was kinetically monitored by qPCR. Lines of best fit were determined by linear regression. The slope of each line is shown, which represents the rate of DNA uptake. The derived slope values serve as a functional correlate for the frequency of T4aP dynamics and are presented as bar graphs in **Fig. 3B**. (**E**) The frequency of dynamic activity in **B** was plotted against the rate of DNA uptake in **C** at the indicated doses of IPTG to assess whether these properties are directly correlated. Linear regression analysis was performed and the R^2^ value is shown on the plot. All data are from 3 independent biological replicates and shown as the mean ± SD. Statistical comparisons were made by one-way ANOVA with Tukey’s multiple comparison test of the log-transformed data. ns = no significance, * = *p* < 0.05, ** = *p* < 0.01, *** = *p* < 0.001. Statistical identifiers directly above bars represent comparisons to the parent.

**Table S1.**
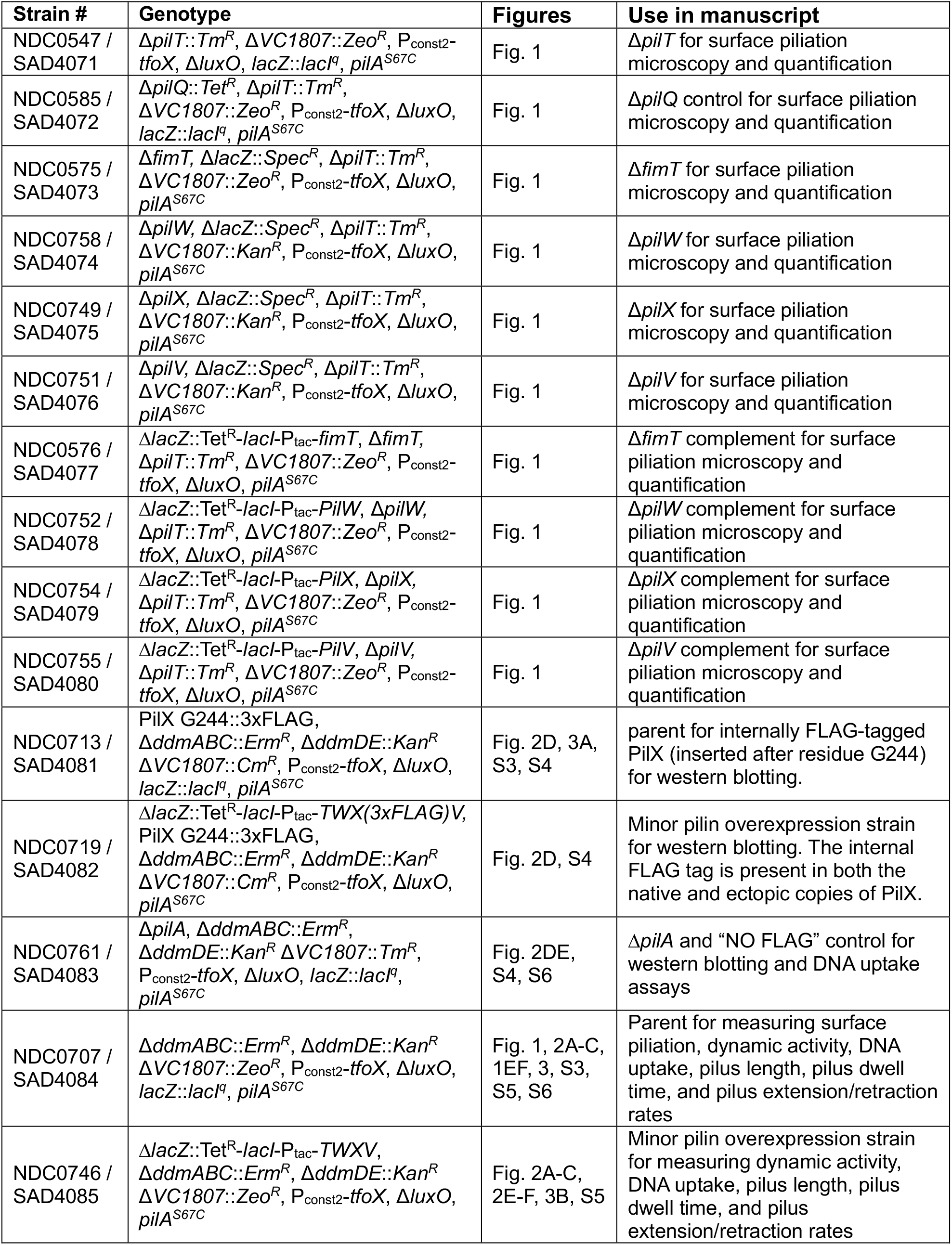
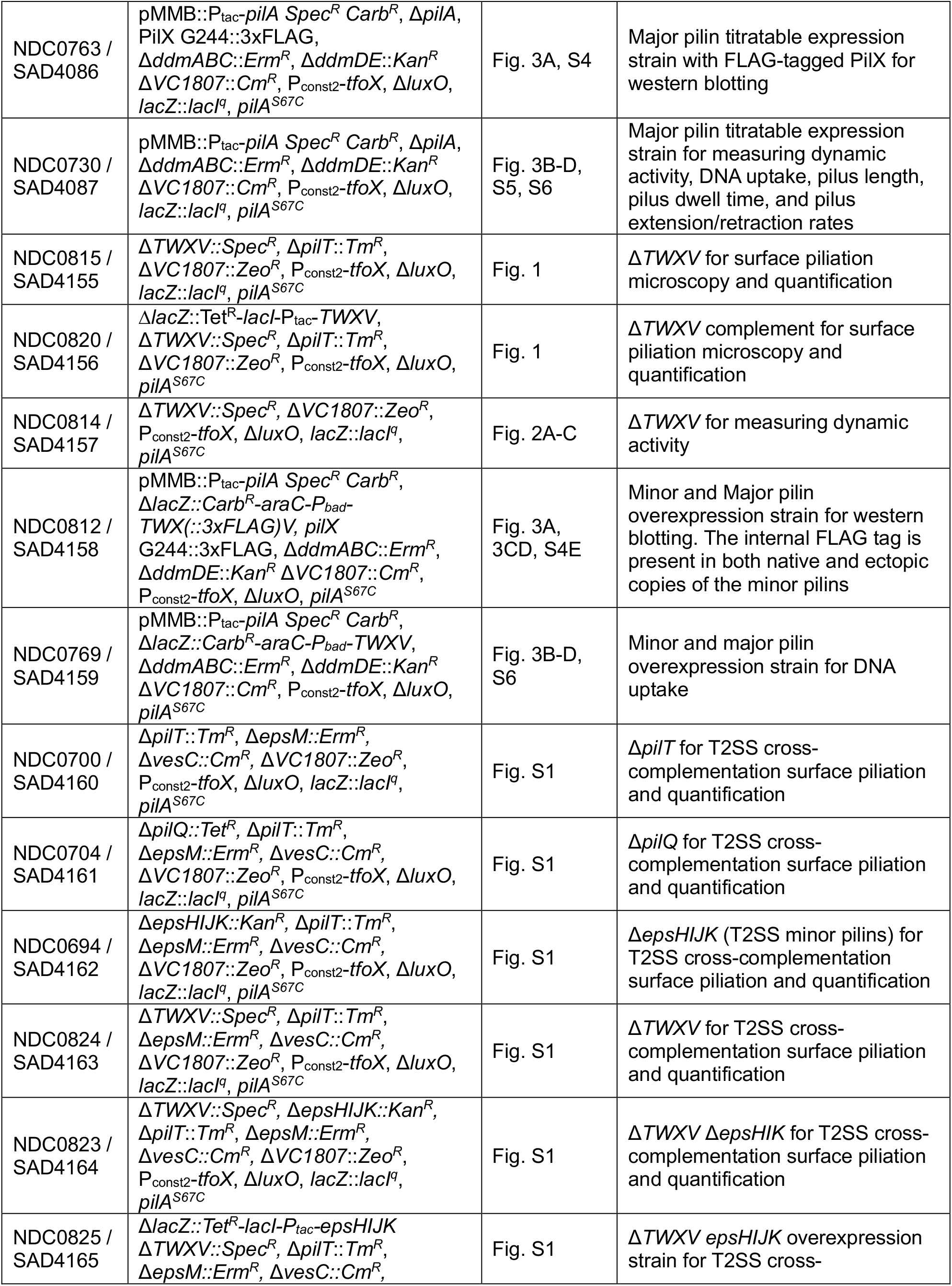
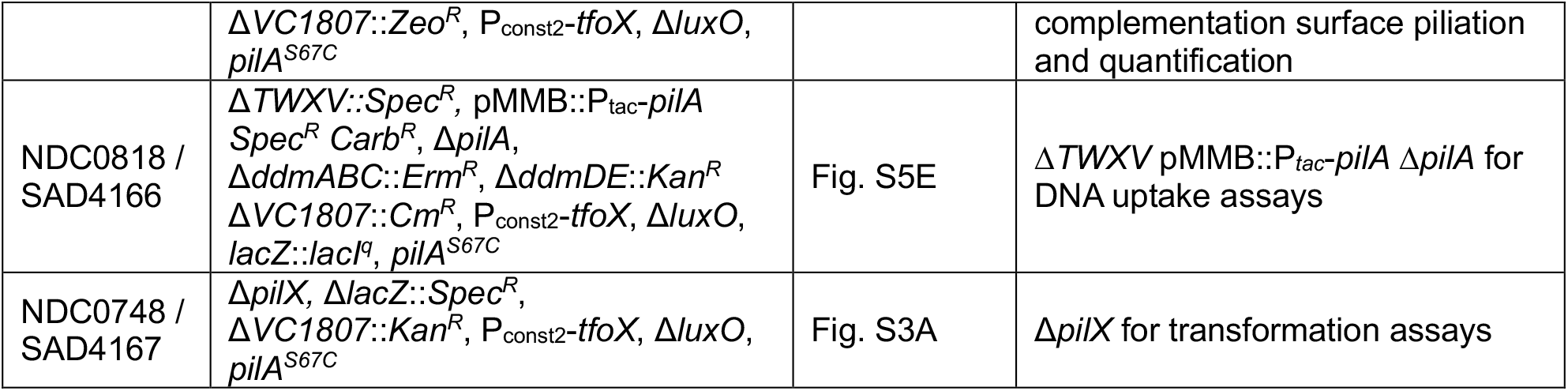
Strains used in this manuscript.

**Table S2.**
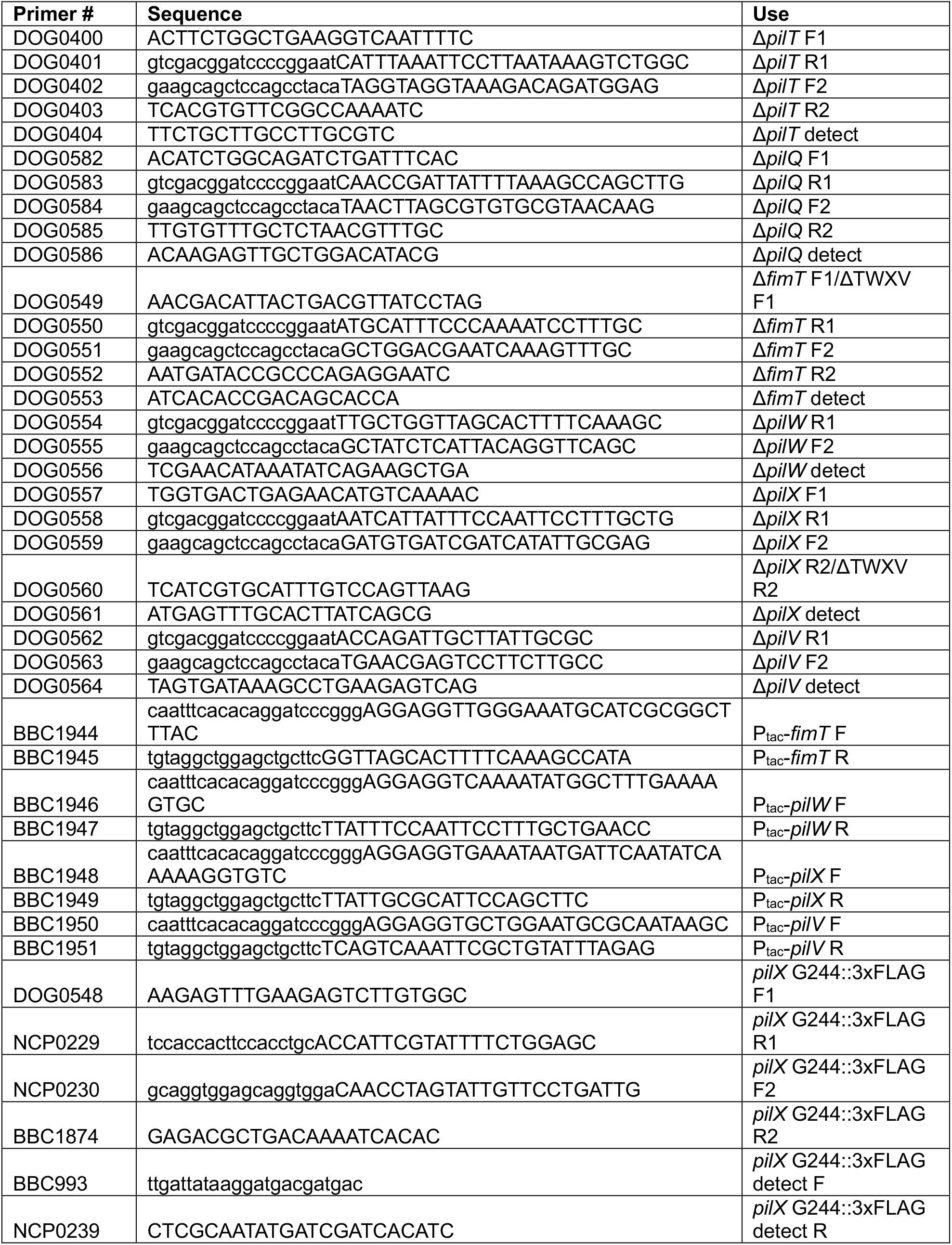
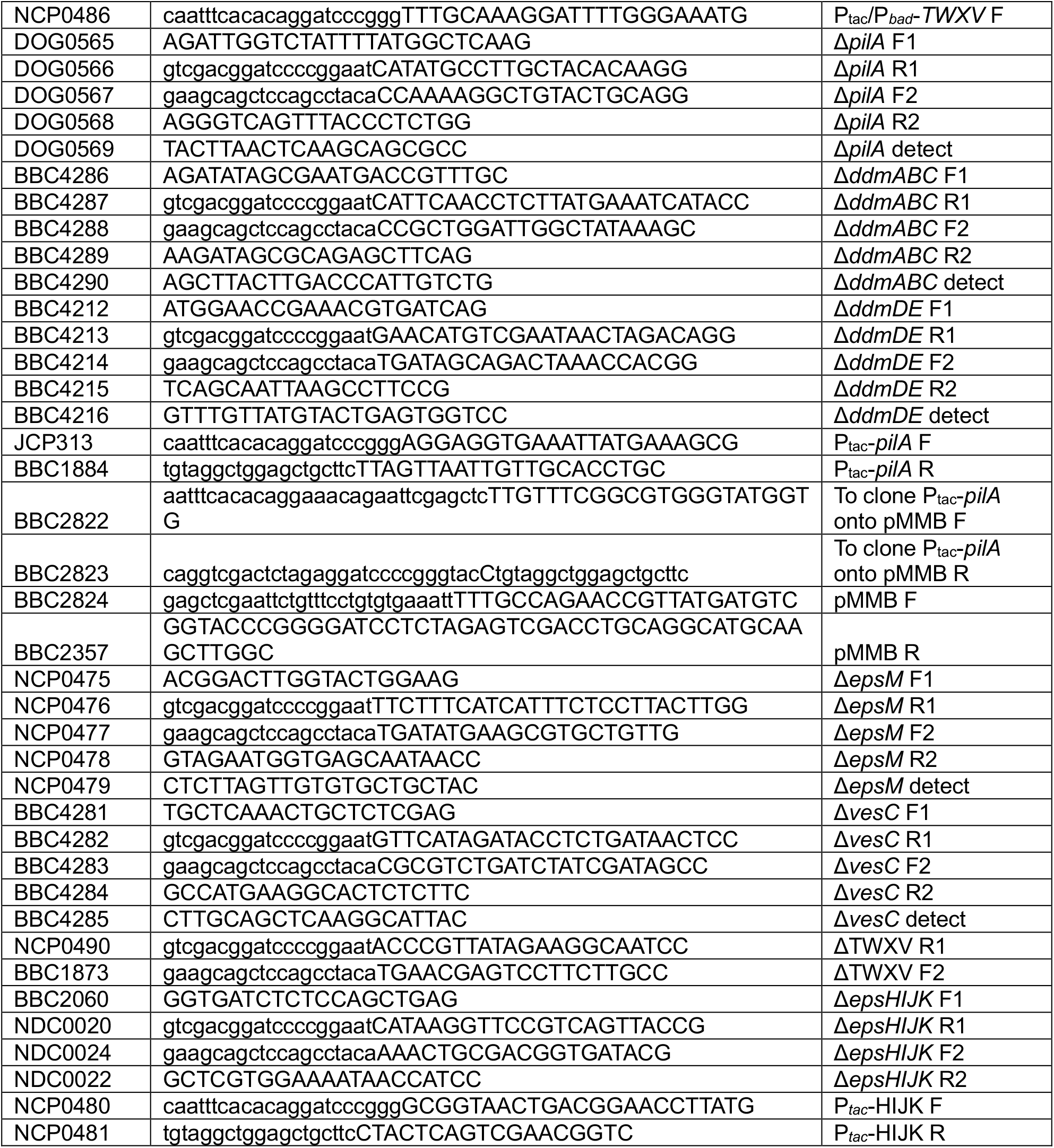
Primers used in this study.

## Notes

### Competing Interest Statement

The authors have declared no competing interest.

### Summary of Updates

Numerous updates to the figures and text that all provide more evidence to support the conclusions of the original submission.

